# Cellular Adaptive Response and Regulation of HIF after Low Dose Gamma-radiation Exposure

**DOI:** 10.1101/316943

**Authors:** Nitin Motilal Gandhi

## Abstract

**Purpose:** Cellular damage due to low dose of γ-radiation (≤0.1 Gy) is generally extrapolated from observing the effects at higher doses. These estimations are not accurate. This has led to uncertainties while assessing the radiation risk factors at low doses. Although there are reports on the radiation induced adaptive response, the mechanism of action is not fully elucidated, leading to the uncertainties. One of the outcomes of low dose radiation exposure is believed to be adaptive response. The mechanism of adaptive response is not fully understood. Therefore, the study was undertaken to understand the role of hypoxia inducible factor (HIF) on radiation induced adaptive response.

**Materials and methods:** MCF-7 cells pre-exposed to low dose γ-radiation (0.1 Gy; Priming dose) were exposed to 2 Gy (challenging dose) 8 hrs after the priming dose and studied for the adaptive response. Cell death was measured by MTT assay, and apoptosis was measured by FACS analysis. DNA damage was measures by alkaline comet assay. HIF transcription activity was assayed using transiently transfected plasmid having HIF consensus sequence and luciferase as the reporter gene.

**Results:** Cells when exposed to 0.1 Gy priming dose 8 hrs prior to the higher dose (2 Gy; Challenging dose) results in lower amount of radiation induced damages compared to the cells exposed to 2 Gy alone. Cobalt chloride treatment in place of priming dose also results in the protection to cells when exposed to challenging dose. There was up-regulation of HIF activity when cells were exposed to priming dose, indicating the role of HIF in radiation induced response.

**Conclusion:** Results indicate the γ-radiation induced adaptive response. One of the mechanism proposed is up-regulation of HIF after low dose exposure, which protects the cells from damages when they are exposed to challenging dose of 2 Gy radiation dose.

## Introduction

The widespread use of medical diagnostic imaging and the fear it has generated has highlighted the need to determine the effects of radiation at low doses (≤ 0.1 Gy). There are uncertainties while measuring the biological effects of radiation in this dose range. (Shore et al. 2017) It is widely accepted that there is no threshold for damaging effect of radiation on living organisms. This is termed as “Linear-no-threshold” (LNT) model. Most of the radiological guidelines are based on this model. There are alternate models put forward like threshold model and adaptive response (Feinendegen et al. 2013; Mullenders et al. 2009). The conundrum is not resolved as all the three models are supported in various studies (Morgan and Bair 2013). These uncertainties in studying the radiobiological effects at low doses are due to the difficulties in assaying radiological damages at low doses accurately. Most of the current knowledge on effects of low dose is based on the atomic bomb survivors by measuring the increase in cancer incidence compared to control population. However, there are several factors which can lead to skewed estimates in population exposed to low dose radiation (Little et al. 2014; Sasaki et al. 2014), as the carcinogenesis being a multi-model disease, it is difficult to estimate the increase in cancer incidence assigned to any one parameter. The use of LNT model results in the stringent conditions to determine the permissible levels of radiation exposure.

It is well known that ionizing radiation damages cellular system through generation of free radicals by breakdown of the water molecule, which is abundant in the cell. These free radicals when scavenged in the cells generate reactive oxygen species (ROS). ROS under these conditions functions as an important physiological regulator of HIF (Finkle T 2012) HIF is a transcription factor which is a heterodimeric protein composed of HIF-1(α) and HIF-1(β) subunit. HIF-1(α) is constantly under degradation pathway in response to the oxygen levels; HIF-1(β) is constitutively expressed. HIF binds to HRE (hypoxia response elements) on the consensus sequence of DNA and regulates the corresponding gene. In the presence of oxygen, the proline residue at 402 and 564 of HIF-1(α) is hydroxylated by prolyl-hydroylase domain (PHD), the hydroxylated protein gets attached to von Hippel-Lindau protein, and is broken down through ubquitination and ptoteosomal degradation. HIF activation is known to provide the increased survival in the cells exposed to ionizing radiation. (Semenza 2012a)

HIF can also be experimentally stabilized through the pharmacological intervention. The function of PHD requires α-ketoglutarate as co-substrate and ferrous ions in their catalytic site. Therefore, iron chelator like desferrioxamine (Woo et al. 2003), and CoCl_2_ can be used to stabilize HIF under normoxic conditions (Yuan et al. 2003)

The work reported here describes the adaptive response in the cells which pre-exposed to the low dose radiation prior to the challenging dose compared to the radiation damage in the cells exposed to the challenging dose alone. One of the mechanism of adaptive response is proposed is due to the activation of HIF on γ-radiation exposure. This stems due to the observation that the CoCl_2_ treatment in place of priming dose can also results in reducing the cellular toxicity by challenging dose.

## Materials and Methods

### Chemicals

Dulbecco’s Modified Eagle Medium (DMEM), Fetal Calf Serum, antibiotic solution, trypsin-EDTA was from Hi media, Mumbai India. Anti-HIF antibody, Annexin V/Pi kit was from BD bioscience. Propidium iodide, Radioimmunoprecipitation (RIPA) buffer, was obtained from Sigma-Aldrich Corp. (St. Louis MO). Lipofectamine3000 was from Invitrogen. Other chemicals used were of analytical grade obtained from local suppliers.

### Cell culture

Human breast cancer cell line MCF-7 was grown in vitro in DMEM with 10% v/v fetal bovine serum. 50 units/ml penicillin and 50 μg/ml streptomycin were added to the media. The cells were grown in CO_2_ incubator set at 5% CO_2_ and 95% air, at 37 °C with relative humidity greater than 85%.

### Radiation exposure

The cells were subjected to γ-irradiation at room temperature (dose rate: 1 Gy/min) using ^60^Co gamma teletherapy machine (Bhabhatron-II, Panacea Medical Technologies, Bangalore, India).

### Cell viability assays

To determine the toxicity of γ-radiation exposure on MCF-7 cells viability assay using MTT assay were conducted. CCK-8 reagent (Sigma) was used to assay the viability of cells treated with CoCl_2_.

### Single cell gel electrophoresis (Comet assay)

The basic alkaline technique described by Singh et al. (2000) was followed with some modifications (Gade and Gandhi 2015) Cell suspension containing 10,000 cells were mixed with 200 μl of 0.7% low melting agarose at 37 °C in a microfuge tube and spread on a fully frosted microscopic slide. The slide was covered with a cover slip and left on ice-cold surface for 2 min. After gelling, cover slip was gently removed. The cells were lysed by dipping the slides into lysing solution (100 mM Na-EDTA, 10 mM Tris, 2.5 M NaCl, 1% Triton X-100 and 10 % DMSO, pH 10) for 1 hr at 4 °C. The slides were rinsed free of salt and detergent in a buffer (1 mM Na-EDTA, 300 mM NaOH, pH>13) and subsequently submerged in a horizontal gel-electrophoresis apparatus by adding fresh buffer, and left in the buffer for 20 min to allow the unwinding of the DNA and the expression of alkali labile damage. Then, an electric field was applied (300 mA; 25 mV) for 20 min to draw negatively charged DNA towards the anode. After electrophoresis, the slides were washed twice for 5 min in a neutralizing buffer (0.4 M Tris, pH 7.4) and stained with 75 μl of propidium iodide (20 μg/ml), and visualized using a fluorescent microscope (Carl Ziess Axioskop) attached with high-performance Carl-Zeiss camera. Fifty cells/slide were captured. The quantification of the DNA strand breaks of images was done using the CASP software (Konca et al. 2000) by which Olive tail moment (OTM) was obtained. OTM is measure of DNA damage which incorporates the percent DNA in tail and tail length.

### Apoptosis assay

Annexin V-FITC/PI dual staining was used for estimating apoptotic cells. Cells were labelled with 5 μl of Annexin V-FITC and 2 μl of propidium iodide (PI, 100 μg/ml) using apoptosis detection kit (BD pharmingen) as per the manufacturer’s protocol. Within one hour the stained cells were acquired in a Partec CyFlow Space™ flow cytometer. Percentage apoptotic cells are determined from the scatter plots using FloJo software.

### Cell-based reporter assay

Cell-based luciferase assay was used to study the transcription activity of HIF. A luciferase reporter gene under the control of hypoxia response elements from the erythropoietin gene (pTK-HRE3-luc; obtained from Prof. S. McKnight, University of Texas, USA) was employed to monitor the HIF activity (Tian et al. 1997) MCF-7 cells were transiently transfected using lipofectamine3000 (Invitrogen) following manufacturer’s protocol by the plasmid and then were exposed to the γ-radiation in different conditions as shown in figure legend. The transfection was continued for 32 hrs, at that time almost 30-50% transfection efficiency. The cells where then exposed to the radiation at various doses. Cells were then harvested, counted in haemocytometer and then lysed with RIPA buffer containing protease inhibitors; snap-frozen under liquid nitrogen and stored at −80 °C. Before analysis the cells were thawed on ice centrifuged at 10,000 g for 5 minutes and the supernatant was analyzed for luciferase enzyme using Sigma-Aldrich (Luc-1) kit following manufacturer’s protocol. The luciferase assay was normalized by equal number of cells taken for assay. The detailed experimental protocol followed is given in individual figure legend.

### Statistical analysis

All data are expressed as the mean ± SEM from three independent experiments. Statistical analyses were performed by the Student’s *t* test, and significance was assumed if *P*<0.05. Statistical significance was measured using Mann-Whitney test while assuming the significance in comet assay results.

## Results

### Optimizing the concentration of Cobalt chloride treatment

Cobalt chloride treatment is known to stabilize HIF-1(α) under normoxic conditions. In order to select the nontoxic but an effective concentration of CoCl_2_; toxicity of CoCl_2_ was determined by exposing the MCF-7 cells under various concentrations of CoCl_2_ for 48 h (Figure 1). CCK-8 assay results indicate the dose dependent toxicity of CoCl_2_ on the viability of MCF-7 cells. At 100 μM and 200 μM there is 8.72% and 17.4% of the toxicity to the cells respectively. Stabilization of HIF after various doses of CoCl_2_ was observed, by western blot analysis (results not shown) In subsequent studies, the concentration 100 μM and 200 μM of CoCl_2_ was used as indicated in the individual figure legend.

**Figure 1.**
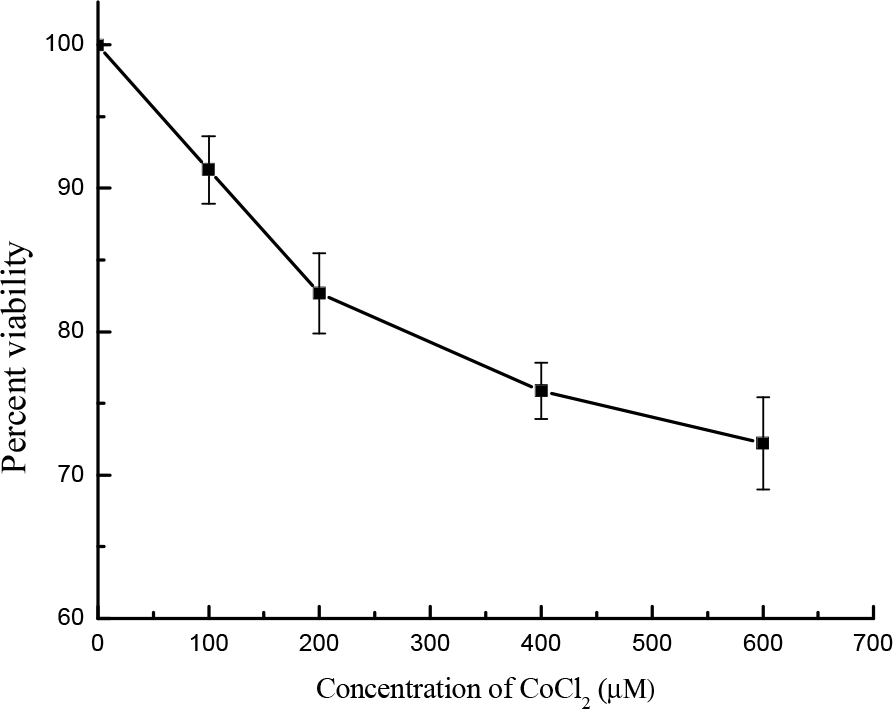
Toxicity of CoCl_2_ on MCF-7 cells. MCF-7 cells were grown in 96 wells plate with different concentration of CoCl_2_ in triplicate. After 48 hours, viability was determined using CCK-8 kit (Sigma). Figure shows the percent viability of MCF-7 cells at different concentration of CoCl_2_ compared to control (without CoCl_2_)

### Adaptive response using Cell viability (MTT) assay

MCF-7 cells grown in 96 well culture plates until a confluence of approximately 70%. The different groups were treated as indicated in the Figure 1 in triplicate. After the incubation for 48 h MTT assay was performed. OD570 of control cells were taken as 100 % and then the absorbance of the other groups was factored. Priming dose alone (0.1 Gy) resulted in slight but statistically significant increase in proliferation. Results depicted in Figure 2 indicate that cells when exposed to 2 Gy γ-radiation alone resulted in significantly higher decrease in viability. Priming dose of 0.1 Gy 8 h prior to the challenging dose of 2 Gy rescued the cells from challenging dose induced decrease in viability. CoCl_2_ treatment in place of priming dose also resulted in the decrease in damage when challenged with 2 Gy γ-radiation.

**Figure 2.**
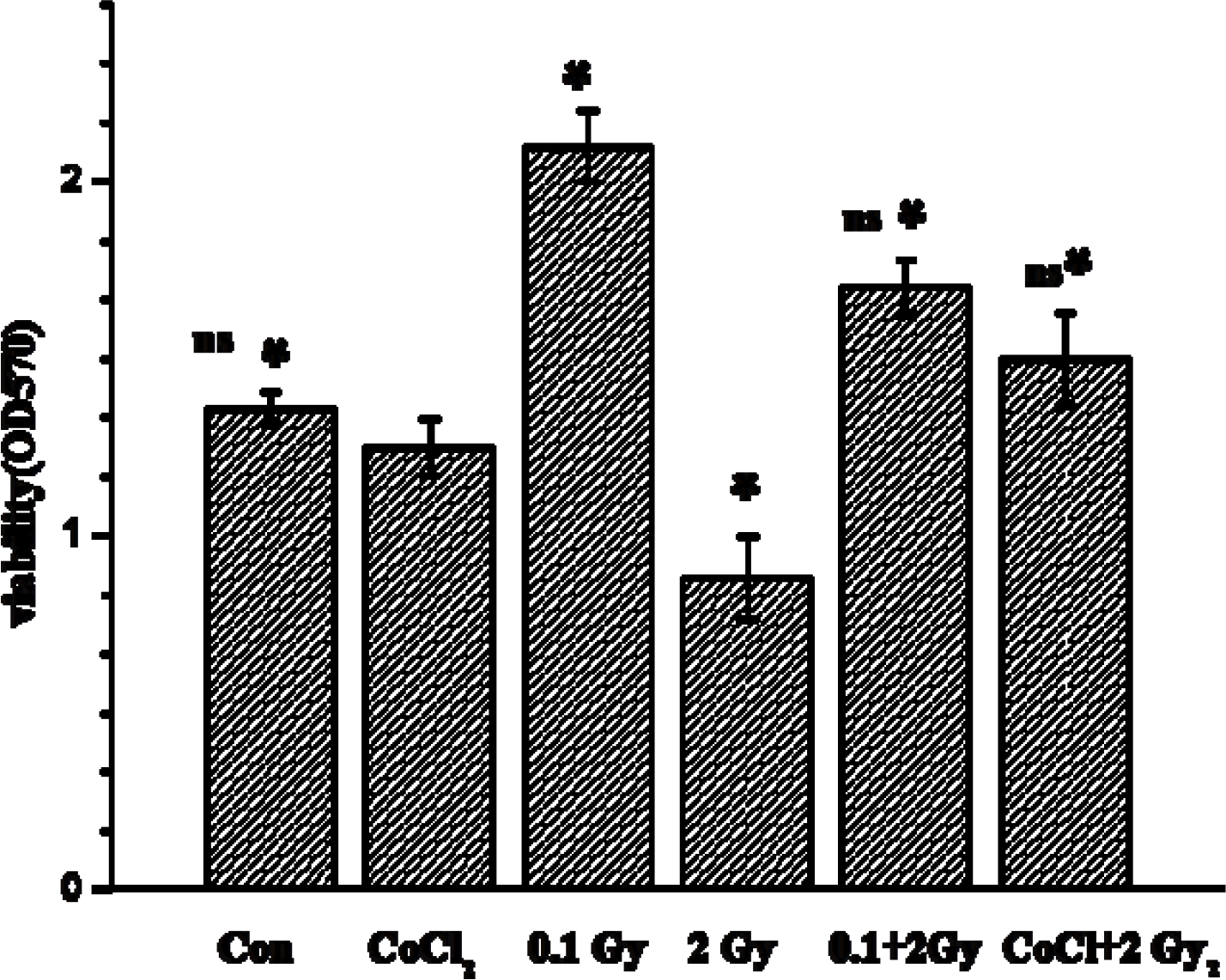
Effect of priming dose on viability of cells when challenged with higher dose assayed by MTT assay MCF-7 cells were first primed with 0.1 Gy of γ-radiation, with one set cells were treated with 100 μM of CoCl_2_ at time T=0, followed by exposure to 2 Gy, after 8 h. MTT assay was performed 48 h post challenging dose. All the sets were done in triplicate. (* p<0.05; * compared to 2 Gy alone and ns= not significant compared to control)

### Measurement of DNA damage using alkaline comet assay

MCF-7 cells grown in 6 well plates were treated with various conditions in triplicate as described in the legend of Figure 3. Immediately after the challenging dose of 2 Gy of γ-radiation, cells were harvested; aliquot of cells (approx.10000 cells) were taken into low melting agarose and subjected to alkaline comet assay. Results indicate that cells when exposed to 2 Gy γ-radiation alone resulted in a significantly more DNA damage as measured as OTM (Fig. 3) Priming dose of 0.1 Gy 8 h prior to the challenging dose of 2 Gy rescued the cells from undergoing the challenging dose induced DNA damages. CoCl_2_ treatment in place of priming dose also resulted in the decrease in 2 Gy γ-radiation induced DNA damages.

**Figure 3.**
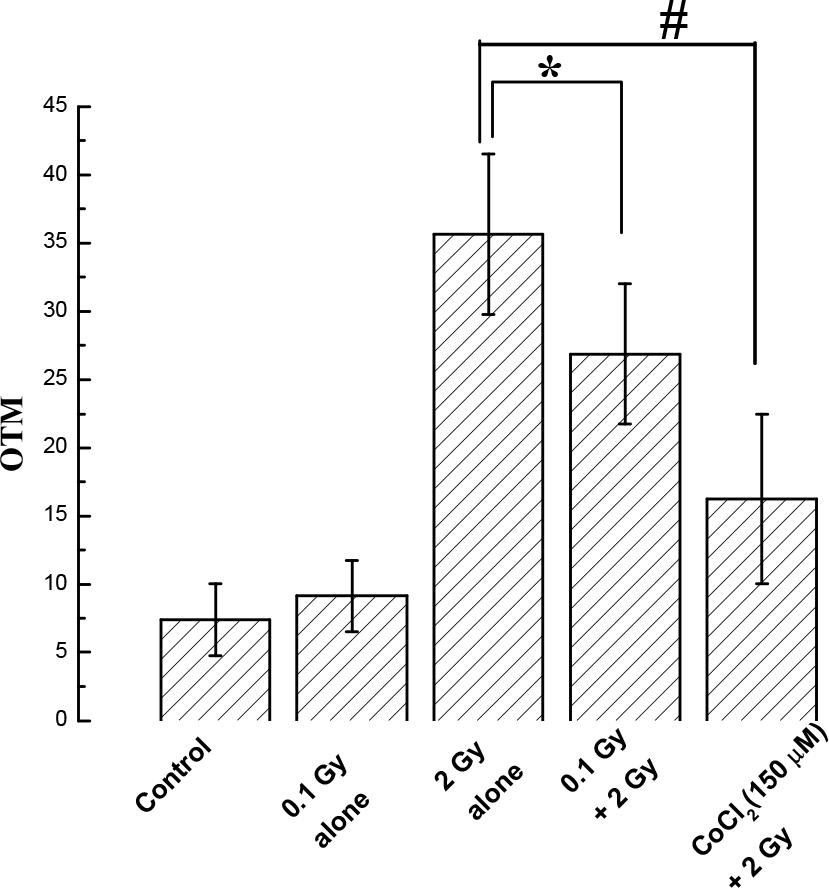
Effect of priming dose on challenging dose induced DNA damage assayed by alkaline comet assay MCF-7 cells were first exposed to the priming dose of 0.1 Gy, after 8 h those cells were exposed to 2 Gy γ-radiation and immediately assayed for DNA strand breaks by comet assay. In one group, cells were treated with 200 μM of CoCl_2_ 7 h before exposing to 2 Gy γ- radiation. All the sets were repeated three times. Statistical significance was measured using Mann-Whitney test (*p<0.05; #p<0.001)

### Measurement of γ-radiation induced Apoptosis

MCF-7 cells were grown in six well culture plates overnight till a confluence of approximately 70% was reached. Different groups were treated in triplicate; after incubation for 48 h cells were harvested, stained by Annexin V/PI and analyzed by FACS. Results indicate that cells when exposed to 2 Gy γ-radiation alone resulted in a significantly higher percentage of apoptosis after 48 h. compared to sham irradiated group. Priming dose of 0.1 Gy 8 h prior to the challenging dose of 2 Gy rescued the cells from undergoing the challenging dose induced apoptosis. (Figure 4) CoCl_2_ treatment in place of priming dose also resulted in the decrease in radiation induced percent apoptosis when exposed to 2 Gy challenging dose. There was no change in percent apoptosis in cells exposed to either 0.1 Gy radiation or cobalt chloride treatment alone (results not shown)

**Figure 4.**
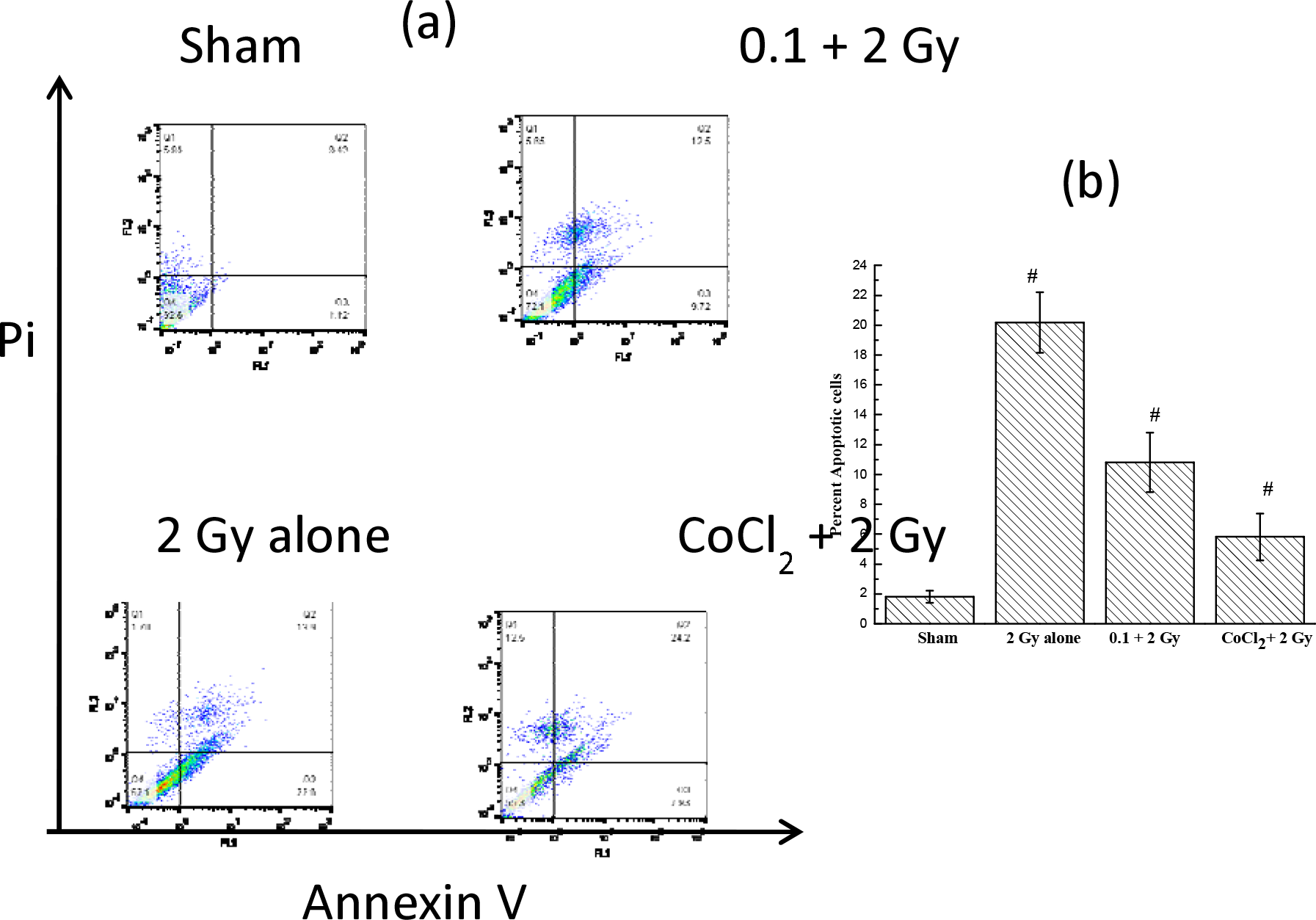
Effect of priming dose on apoptotic cell death after exposure to challenging dose of 2 Gy MCF-7 cells were first primed with 0.1 Gy of priming dose. After 8 h of incubation cells were exposed to 2 Gy γ-radiation. After challenging dose cells were further incubated for 48 h, harvested and stained with FITC-Annexin V/Pi and analyzed by FACS. One group was treated with CoCl_2_ (200 μM) in place of priming dose. (#p<0.05 with student’s t test compared to control)

### Modulation of γ-radiation induced HIF transcription activity

To increase the sensitivity to measure the HIF regulation at lower dose, transiently transfected MCF-7 cells with p-HRE3-tk-luc were exposed to various doses of γ-radiation. The plasmid contains luciferase gene under control of HIF consensus signal. Therefore, the luciferase signal is directly proportional to the transcription activity of HIF. After 8 h the cells were lysed and analyzed for luciferase activity. There was dose dependent increase in luciferase signal, there was significantly increase in luciferase activity at 0.1 Gy compared to un-irradiated control. The signal saturated at 0.5 Gy as there was no statistically significant increase in luciferase activity between 0.5 Gy and 2 Gy. CoCl_2_ treatment alone (used as positive control) gave very strong increase in luciferase activity (Figure 5).

**Figure 5.**
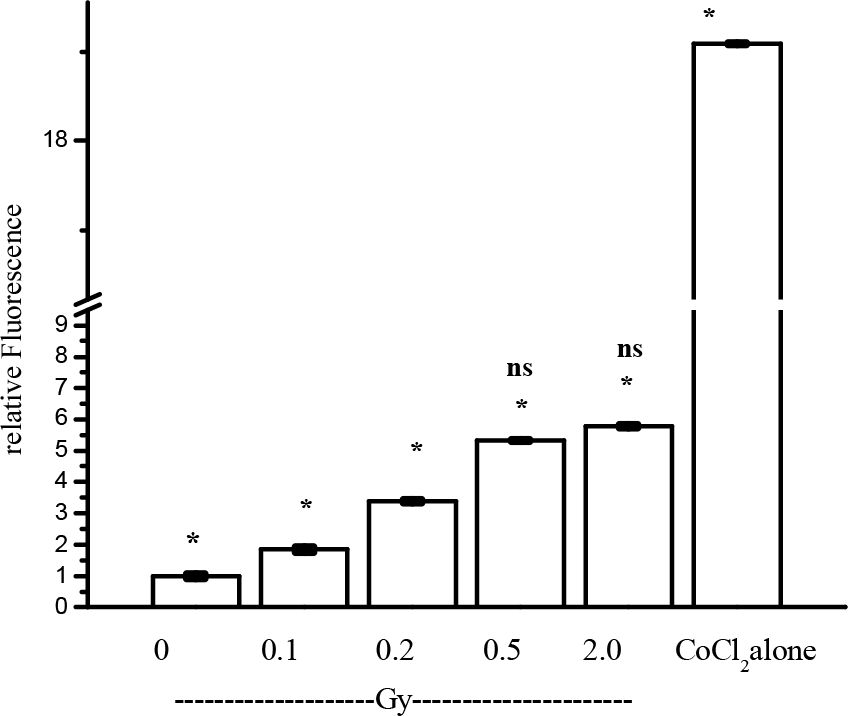
Dose dependant induction of transcription activity.MCF-7 cells were plated on 96 well plates 24 h prior to transfection. After washing out the media, the cells were incubated with 50 ng of p-HRE3-tk-luc plasmid and lipofectamine reagent in media without FCS. After 4 h of incubation 10% FCS was added the cells were further incubated for 16 h. The cells were then exposed to different doses of γ-radiation and incubated for 8 h. In one set CoCl2 (200 μM) was added as positive control. All the experiment set were done in triplicate. At the end of the incubation the cells were harvested, counted for the cell number and were snap-frozen in liquid nitrogen, and then stored at −80 ±C. Luciferase assay was performed using LUC-1 kit (Sigma). Significance was assumed if p<0.05. (* compared to non-irradiated; 0 Gy control and, ns= not significant, compared to each other.)

### Discussion

In the present study, the adaptive response in MCF-7 cells is described. Viability, apoptosis and DNA damage outcomes have suggested that there is a significant reduction of the radiation induced damages to cells pre-exposed to 0.1 Gy γ-radiation. Further, the results also suggest that when 0.1 Gy priming dose was replaced with CoCl_2_ treatment resulted in reduction of damages after the challenging dose.

The chemical toxicity of Cobalt chloride treatment is minimal as shown in Figure 1. However, there is statistically significant adaptive response when priming dose if replaced with Cobalt chloride treatment. The toxicity induced by cobalt chloride could not be attributable to the adaptive response as any chemical toxicity induced adaptive response induction takes longer time duration (if at all) cobalt chloride treatment is given for 8 h, time not sufficient to make the global effect of chemical stress induced adaptive response. These results indicate that CoCl_2_ treatment (and thus HIF stabilization) can mimic the effects of priming dose. This finding was corroborated as when transiently transfected cells with p-HRE-tk-luc plasmid was treated with different radiation doses; there was dose dependent increase in HIF transcription activity. Further, at 0.1 Gy there was significant increase in transcription activity of HIF indicating the activation of HIF. In spite of several reports which describe the γ-radiation induced adaptive response in cells; these observations could not translate to the reduction of stringent radiological guidelines. One of the reasons may be due to the lack of understanding in molecular mechanism which can explain the adaptive response. Lall et al. (2014) have shown the adaptive response in human fibroblasts is due to the increased in glucose metabolism. Their results also suggest that at higher doses of γ-radiation p53 gets activated which in turn shuts off the HIF stabilization, contrary to findings reported here, the difference of observations is assumed to stem from the cell type.

There are reports on studies on adaptive response in MCF-7 cells (Min et al. 2005; and Suwimol et al. 2008) where authors have shown the activation of MnSOD as one of the mechanism of adaptive response. Alexander et al. (2003) have demonstrated that NF-κB regulates the expression of MnSOD, which in turn increases the expression of genes that participate in radiation-induced adaptive response in MCF-7 cells.

Zhou et al. (2013) have shown that the combination of radiation and hypoxia could promote the glucose metabolism. Further, HIF-1(α) might inhibit the promoting effect of radiation on glycolysis in hypoxic MCF-7 cells by regulating the glucose metabolism. MCF-7 has wild type p53 status. There is the direct correlation with p53 and HIF therefore it was essential to study the radiation induced modulation of HIF in MCF-7 cells (Murley et al. 2017).

One of the observation that exposing the cells, when exposed to 0.1 Gy alone resulted in statistically significant increase in cell proliferation compare to non-irradiated control cells. This could be additional evidence that ionizing radiation can enhance the proliferation of cells and promote long term resistance to multiple cytotoxic stresses (Anupama and Rajgopal 2013) Ionizing radiation exposure cause the disruption in the cellular functions or cause death to the cells depending on the dose of radiation. Ionizing radiation damages the cellular functions by generating free radicals and ROS through breakdown of water molecule. Radiation exposure results in cellular damages to DNA, protein and lipids. Those damages result in changes in biochemical and molecular signalling in the surviving fraction of cells (Azzam et al. 2011). One of the factor that get modulate is HIF.HIF is primarily an oxygen sensing transcription factor, under the continuous degradation pathway in the presence of oxygen; however, under the hypoxic conditions HIF is stabilized and regulates almost 800 genes (Semenza 2012b) It is also known that hypoxia is not the only cause of stabilization of HIF. Even in the presence of oxygen HIF can get stabilize through the effect of decrease in Fe2+ (Bianchi et al. 1999) or the presence of intermediate of TCA cycle like alpha-ketoglutarate (Pan et al 2007) The significant increase in HIF transcription activity at 0.1 Gy radiation was observed. As exposure to γ-radiation results in activation of MAP kinase pathway through EGFR signalling (Michael et al 2000) Further, Mylonis et al. (2006) have shown that MAPK-dependent phosphorylation of HIF is required for its efficient accumulation inside the nucleus. Taken together this could explain the increase in the transcription activity due to enhance in nuclear translocation of HIF-1(α).

The background radiation dose was greater than eight times when life was evolved, compared to the present day (Karam and Leslie 1999). Which could have resulted in evolution of efficient DNA repair system, which is evolutionary conserved till the present day this could help the humans to repair the low dose of radiation induced damages efficiently. Therefore, we have to review the notion that even a very low dose of radiation is harmful to the life. Thus, the debate regarding LNT model needs to be reviewed.

Taken together, based on the findings presented here, it is proposed that the low dose of γ- radiation results in the increased transcription activity of HIF and subsequent activation of survival pathways. Therefore, when the cells were primed with low doses of radiation, it resulted in up-regulation of survival machinery protecting the cells when they were challenged with higher doses of radiation.

## Acknowledgements

This work was supported by Department of Atomic Energy, Government of India. Author will like to thank Dr. PV Babu for the critical review of the manuscript. Dr. Steve Mcknight for generously providing the plasmid. Ms. Shruti Gade and Ms. Krishna for helping in some of the experiments. Mr. Prayag Amin for conducting the FACS analysis.

